# Neuronal control of the fingertips is socially configured in touchscreen smartphone users

**DOI:** 10.1101/064485

**Authors:** Myriam Balerna, Arko Ghosh

**Author notes:** Corresponding author: Arko Ghosh, Institute of Neuroinformatics, University of Zurich and 14 ETH Zurich, Winterthurerstr. 190, 8400 Zurich, Switzerland. **Author Contributions:** MB acquired the data, participated in data analysis, and edited this manuscript. AG designed the study, helped in data acquisition, analyzed the data, and drafted this manuscript.

## Abstract

As a common neuroscientific observation, the more a body part is used, the less variable the corresponding computations become. We here report a more complicated scenario concerning the fingertips of smartphone users. We sorted 21-days histories of touchscreen use of 57 volunteers into social and non-social categories. Sensorimotor variability was measured in a laboratory setting by simple button depressions and scalp electrodes (electroencephalogram, EEG). The ms range trial-to-trial variability in button depression was directly proportional to the number of social touches and inversely proportional to non-social touches. Variability of the early tactile somatosensory potentials was also proportional to the number of social touches, but not to non-social touches. The number of Apps and the speed of touchscreen use also reflected this variability. We suggest that smartphone use affects elementary computations even in tasks not involving a phone and that social activities uniquely reconfigure the thumb to touchscreen use.

**Impact Statement:** Unconstrained behavior on a smartphone is a powerful predictor of neuronal functions measured in the laboratory and the details of the smartphone-neuronal association challenges the established ideas of brain plasticity.

## Introduction

Smartphones enable a remarkably broad range of activities. From the perspective of higher cognition, smartphone behavior engages complex computations for decision-making, language, and social interactions. From the perspective of lower-level sensorimotor control, the thumb and the fingertips are repeatedly applied on the touchscreen to essentially either tap or swipe. The observation that even toddlers can easily operate a touchscreen underscores the simplicity of its sensorimotor control (1). According to a series of experiments, a repeated use of the hand in either skillful or simple actions enhances the corresponding representation in the sensorimotor cortex (2–6). Sensorimotor alterations have been observed in trained laboratory monkeys, athletes, Braille readers, and concert string instrument players (3, 5, 7–9). A prominent notion underlying these observations is that the sensorimotor cortex keeps track of the amount of activity generated by the corresponding body part but the exact nature of this tracking is unclear. For instance, in terms of touchscreen use, the cortex may keep track of the number, frequency, and/or behavioral context of touchscreen actions.

In real-world observations, the role of the behavioral context in use-dependent plasticity is difficult to establish, partly because of a poor quantification of human actions. For instance, it is common to assess the extent of deliberate practice in elite musicians by using questionnaires (6, 10, 11). Such qualitative approaches do not provide a measure of the amount of activity nor do they capture the activity context. Under well-controlled laboratory conditions, the precise extent of plasticity depends on whether the sensory information presented at the fingertip is used towards a behavioral task or not (4). In general, the cortical plasticity can be modulated by artificially stimulating neuromodulators, such as dopamine or serotonin, that are naturally released according to the behavioral relevance (12). Social behavior strongly engages such neuromodulators and the touchscreen smartphone is prominently used towards social activities (13–15). Therefore, the use-dependent configuration of fingertips in touchscreen users might not be a simple function of sensorimotor activity (16). In particular, touchscreen touches used towards social activities may be distinctly weighted towards use-dependent plasticity of the sensorimotor cortex. Social activities are well compartmentalized within specific Apps, allowing us to quantitatively address use-dependent plasticity in distinct behavioral contexts.

In this report, we focused on the elementary property of neuronal variability, or noise, in the sensorimotor system. Substantial theoretical and empirical support exists for the notion that an increased use of a body part reduces the sensorimotor noise (17–21). According to one prominent theory, the brain actively learns to suppress motor variability as if to eliminate unwanted noise, albeit a different theory has been put forward on how the brain may exploit the inherent noise towards learning (18, 22). Sensorimotor variability of the fingertips is diminished with musical practice, by typing on the keyboard, or by deliberately practicing laboratory-designed tasks (18, 23–25). Therefore, a clear-cut prediction would be that the sensorimotor variability of the fingertips is diminished with increased touchscreen use, irrespective of the actions being social or non-social. Alternatively, the complexity, neuromodulation, and the overall significance of social activities may distinctly shape the sensorimotor variability.

To address these possibilities, we performed a multiple regression analysis to assess the relationship between (a) Social App usage in the real world and sensorimotor variability measured in the laboratory, and (b) Non-social App use and sensorimotor variability measured in the laboratory. We also examined other variables that were likely to influence sensorimotor variability. To alleviate the effect of development or aging on our measurements, we restricted the analysis to a young adult population (26). Gender-associated differences exist in sensorimotor processing from the fingertips and in the performance variability of a simple task (27, 28). Therefore, we included a dummy variable representing the gender of participants in the regression analysis. Since an accurate control of motor timing is important for rapid actions, fast touchscreen operators may develop a more precise sensorimotor system (29). Therefore, a typical rate of touchscreen touches was added as an explanatory variable. Finally, practicing motor skills in various contexts leads to better performance in a previously not experienced context (30). Since each App on the phone is associated with a distinct context, we quantified the number of Apps in use as an explanatory variable. In summary, type of touchscreen activity (social or non-social), the gender, a typical rate of touchscreen activity, and the number of Apps may all impact sensorimotor computations measured in the laboratory. Incorporating these factors in a single regression model allowed us to address if and how they are separately weighing in on the sensorimotor variability.

## Results

### Basic features of touchscreen use

We quantified touchscreen use for a period of 21 d in a young adult population using a custom-designed software operating in the background to record every touchscreen event and the App targeted by the event. Social activity generated on the touchscreen was sorted based on the App in use. We considered Apps that primarily enabled the communication of personal messages or opinions to a circle of friends or acquaintances as “Social”, and Apps that did not fulfill these functions as “Non-social” (for a sample of Social and Non-social Apps in the database see ***Supplementary List 1***). The usage statistics were as follows: the volunteers touched the screen from 1540.3 (20th percentile) to 5562.3 (80th percentile) times per day, and generated between 429.1 (20th percentile) and 2486.9 (80th percentile) touches per day on the Social Apps. Importantly, the number of social touches was only partly correlated with the number of non-social touches [variables Log10 normalized, R^2^ = 0.29, *f* (1,55) = 22, *p* = 1.9 × 10^−6^, robust linear regression]. Furthermore, volunteers ranked the fingers used according to their preference. Confirming previous findings for smartphone usage, the thumb was ranked by 73% of the users as most preferred on the touchscreen; 16% preferred the index finger; and 10% preferentially used both the thumb and the index finger (16, 31). Remarkably, only one user preferred their middle finger to all the other fingers.

### Motor variability of the thumb, but not of the middle finger, is associated with touchscreen use

At the end of the touchscreen recording period, the volunteers performed a simple tactile reaction task in the laboratory where the reaction involved micro switch press-down and release-up actions (***Figure 1a,b***). In theory, the time taken to trigger the press-down action (reaction time) reflects the sensory decision processes, and the time taken to complete the motor act, from pressing down to releasing upwards (movement time), reflects the lower cognitive levels of sensorimotor execution (32–35). The trial-to-trial variability was parametrized using ex-Gaussian fits. Specifically, we estimated the variability of Gaussian curve region lacking very slow actions driven by attention lapses (36, 37). In agreement with the notion that the reaction and movement times reflect distinct neuronal computations, the corresponding variabilities were unrelated to each other [R^2^ = 0.02, *f* (1,53) = 1.1, *p* = 0.299, robust linear regression]. Since we were interested in the low-level sensorimotor variability, we focused on the movement time.

**Figure 1.**
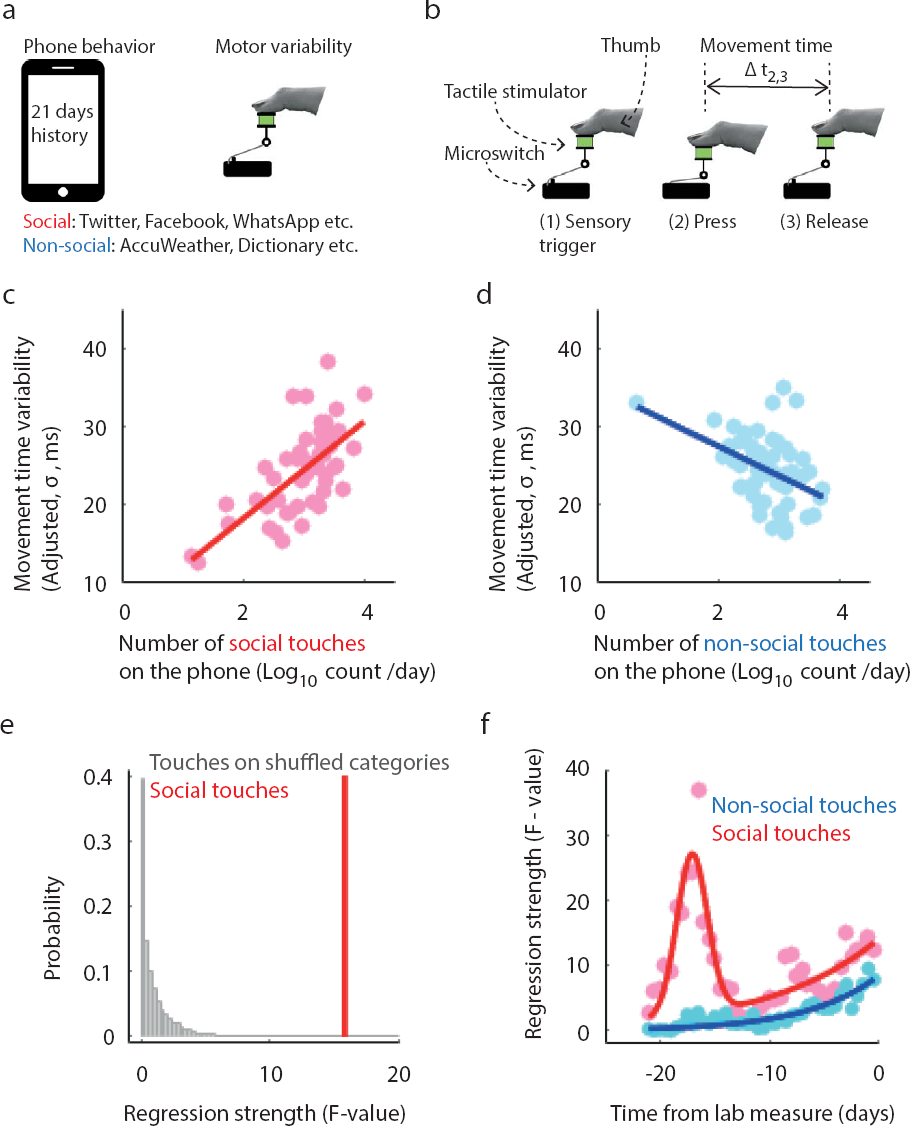
The history of unconstrained touchscreen behavior reflects on the performance of a simple task. (a) Touchscreen activity was recorded for 21 d and followed by laboratory measurements of sensorimotor variability. (b) The task required responding to tactile stimuli by pressing and releasing a micro switch, as fast as possible, with the thumb. (c-d) Adjusted response plots. (c) Movement time variability (σ) was directly proportional to the number of touches generated on the Social Apps (social touches). (d) The movement time variability was inversely proportional to the number of touches generated on the Non-social Apps (non-social touches). (e) The distribution of relationships for randomly categorized Apps (10^4^ iterations) in comparison to the relationship uncovered for social touches. (f) Parsing the touchscreen recordings in 12 h steps (72 h bin) revealed that the relationship involving non-social touches simply decayed as a function of time, whereas the relationship involving social touches showed a more complex pattern. The statistical tests and the details of the fits are reported in the main text.

In our multiple linear regression analysis of movement time variability, we treated the number of daily touches on the Social, Non-social, and Uncategorized Apps (all Log10-normalized), gender (dummy variable, female = 1), typical rate of touchscreen touches, and the number of Apps used during the recording period, as explanatory variables. First, let us elaborate on the thumb use analysis data (the thumb was most preferred for touchscreen interactions). The full regression model was highly significant [R^2^ = 0.45, *f* (6,48) = 6.5, *p* = 4.43 × 10^−5^, robust multiple linear regression; for variation inflation factors see ***Supplementary Figure 1***]. The maximum variation inflation factor was 2.7, indicating that the regression model was not affected by multicollinearity (38). According to the simple prediction of use-dependent reduction in sensorimotor variability, the regression coefficient for social touches was expected to be either zero, suggesting that social actions are not distinctly tracked by the brain, or negative, suggesting that social actions are distinctly tracked but a higher number of social touches leads to lower sensorimotor variability. Contrary to these predictions, we found that higher number of social touches led to increased movement time variability [*t*(1,48) = 3.96, *p* = 0.00024, ***Figure 1c***]. The case for non-social touches was anticipated, with higher number linked with smaller variability [*t*(1,48) = −2.66, *p* = 0.011, ***Figure 1d***]. The same was observed for uncategorized touches [*t*(1,48) = −2.45, *p* = 0.018].

To what extent does the social behavior-movement time variability relationship (***Figure 1c***) depend on App classification? We addressed this by repeating our analysis 10^5^ times using randomly shuffled categories. The relationship uncovered for social touches was well separated from the distribution of relationships obtained by quantifying random category touches (***Figure 1e***). This result further supported the notion that the type of touchscreen behavior determines how neuronal processes responsible for the thumb are configured.

To address whether the touchscreen behavior-movement time variability relationship was specific to the thumb, a subset of volunteers also performed the task with their middle finger (which was rarely indicated as the preferred finger for touchscreen use). We found a strong association between the explanatory variables and movement time variability for the thumb [R^2^ = 0.79, *f* (6,10) = 6.43, *p* = 0.0053, robust linear regression], similarly to data for the full set of volunteers. Importantly, here too the number of social touches was significantly related with movement variability [*t*(1,10) = 2.70, *p* = 0.022, ***Figure 1 – Supplement 1*].**]. However, the results for the middle finger were strikingly different. We found no correlation between the explanatory variables and movement time variability [R^2^ = 0.28, *f* (6,10) = 0.66, *p* = 0.683, robust linear regression]. Moreover, the regression coefficient associated with the number of social touches was non-significant [*t*(1,10) = −0.30, *p* = 0.77, ***Figure 1 – Supplement 1*].**. These results suggested that the putative impact of touchscreen use on movement time variability is specific to the finger that is repeatedly engaged on the touchscreen.

### Social keypad touches distinctly impact on motor variability

In the analyses conducted above, the touchscreen touches consisted of different gestures, i.e., keypad taps, swipes, and pinches. One interesting possibility was that the correlations identified for social touches were driven by the different gestures used for Social Apps. Therefore, we next restricted our analysis to pop-up keypad touches. It is safe to assume that for sensorimotor control, i.e., the degrees of freedom for motor control and visuomotor coordination, keypad touches for Social Apps are the same as the ones for Non-social Apps. The difference concerns the specific content typed. Full regression model based on the keypad touches was significantly related to motor time variability [R^2^ = 0.60, *f* (6,25) = 6.36, *p* = 0.0004, robust linear regression]. We noted that the higher the number of social touches on the keypad, the larger the movement time variability [*t*(1,25) = 3.76, *p* = 0.0009, ***Supplementary Figure 2***]. This suggested that gestures cannot simply account for the distinct imprint of social activities on motor time variability.

### Social and non-social touches show distinct patterns of correlations as a function of time

The continuously recorded touchscreen behavior made prior to the laboratory measurements allowed us to address the question of whether the touchscreen-movement time variability relationship changes as a function of time. Should the relationship be driven by rapid plasticity, then it would simply decay as a function of time. However, if slow mechanisms were operational, then the relationship would peak with older rather than the most recent touchscreen experiences, as if indicating a delayed impact of touchscreen behavior. F-values, describing the relationship strength, revealed a simple decay trend for non-social touches. This was well described (R^2^ = 0.82, ***Figure 1f***) by:

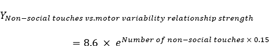

The relationship for social touches was more complicated, consisting of both an initial decay and a strong relationship with older data. This dynamic was well described (R^2^ = 0.81, ***Figure 1f***) by:

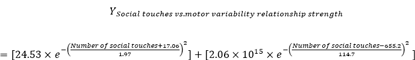

The distinct pattern of time-dependent relationships for social vs. non-social touches suggested that they engage different forms of plasticity.

We also revealed the dynamics of other explanatory variables that were significantly related to touchscreen use recorded over the 21-d period. In brief, as anticipated, variability was smaller with a higher typical rate of touchscreen touches [*t*(1,48) = −5.10, *p* = 5.73 × 10^−6^, ***Figure 1 - Supplement 2*]** and with a larger number of Apps used [*t*(1,48) = -3.29, *p* = 0.002, ***Figure 1 - Supplement 2*]**. Time-dependent dynamics for the typical rate of touchscreen touches indicated slow plasticity but the “number of Apps” variable dynamics indicated both rapid and slow plasticity **(*Figure 1 - Supplement 2*)**. The gender of the user was not significantly associated with the motor time variability [*t*(1,48) = -0.90, *p* = 0.37].

### Social touches distinctly affect the reaction time variability

We opportunistically explored the variability of higher cognitive levels captured by the reaction time. For the reaction time variability, the full regression model was significant but weak [R^2^ = 0.26, *f* (6,49) = 2.86, *p* = 0.02, robust linear regression]. Similarly to the results for movement time variability, we observed that a higher number of social touches was associated with greater reaction time variability [*t*(1,49) = 2.72, *p* = 0.009, ***Supplementary Figure 3***]. The only other explanatory variable that significantly contributed to the regression model was the participant gender, such that the females showed less variability [*t*(1,49) = -3.25, *p* = 0.0002] than the males. Since the reaction and movement times measure different aspects of cognition, taken together, they suggested that the putative impact of social touches is not restrained to the lower-levels of sensorimotor cognition.

### The signal-to-noise ratio of the early somatosensory evoked potentials from the thumb strongly corresponds with touchscreen use

To address the neurophysiological predictions of use-dependent plasticity, we measured the cortical potentials in response to tactile stimulation of the fingertips using electroencephalography (EEG). The EEG signals were noisy at a single trial level and an averaging method across several trials revealed an event-related potential (***Figure 2a***) (39). We used the ratio between the average response and a trial-to-trial deviation from the average as a measure of putative signal-to-noise ratio. Based on the observations from an electrode showing the strongest response (according to the grand average), a distinctive rise in the signal-to-noise ratio was observed, with a peak at 55 ms (latencies are reported from the onset of stimuli, ***Figure 2b***).

**Figure 2.**
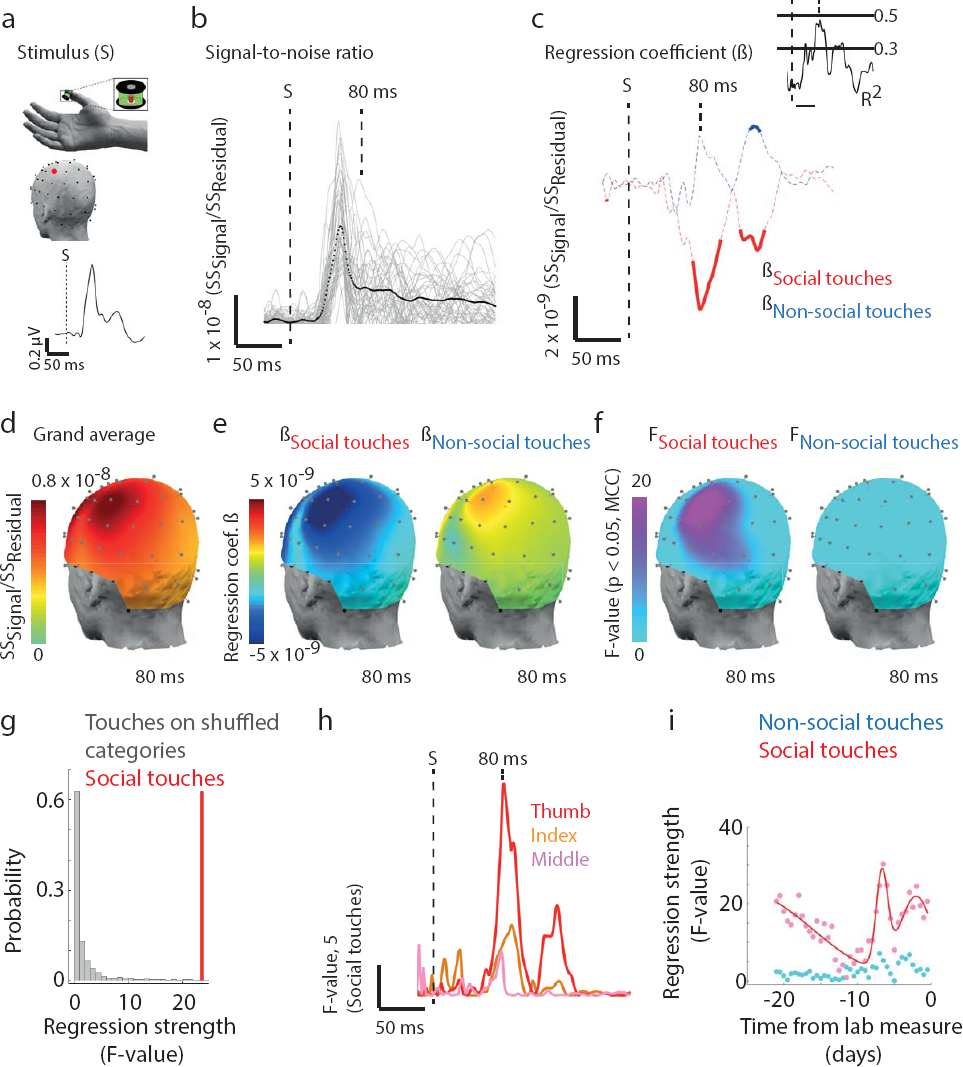
Early cortical somatosensory processing reflects the history of Social App usage. (**a**) We estimated the signal-to-noise ratio in the cortical responses upon a brief tactile stimulus presented to the right thumb tip, the hand was in a resting position during the recording. The head plot shows the electrode location with the best response (red) (**b**) Putative signal-to-noise ratio (SNR) at the electrode (SS, sum of squares). Individual volunteers (gray lines) and population mean (black). (**c**) Event related coefficients with the SNR as dependent variable and touchscreen parameters based on the entire 21 d recordings as explanatory variables. Statistically significant coefficients (thickened lines, p < 0.05, corrected for multiple comparisons, ANOVA). (**d**) Head plot of the population mean of the SNR at a latency of 80 ms. (**e,f**) The event related coefficients and the corresponding statistics at 80 ms. (**g**) At the chosen electrode and at 80 ms, the distribution of the relationship strength based on randomly categorized Apps (10^4^ iterations) in comparison to the relationship uncovered for social touches. (**h**) The relationship with social touches was the strongest for the thumb, followed by the index finger, and, finally, the middle finger. (**i**) Parsing the touchscreen recordings in 12 h steps (72 h bin) revealed that the relationship between social touches and the signal-to-noise ratio evoked from the thumb at 80 ms latency fluctuated in a complex manner through the recording period. The details of the fit is reported in the main text.

We were interested in both the direction and timing of neuronal correlates of touchscreen use. Based on the simplistic prediction of use-dependent plasticity, we anticipated that the more the fingertips are used on the touchscreen (irrespective of the social category of the activity), the larger the signal-to-noise ratio (6, 16, 40). Measurements at different latencies reflect distinct stages of the cortical somatosensory processing, with the potentials between 40 and 100 ms dominated by the primary somatosensory cortex, and those between 100 and 200 ms dominated by the secondary somatosensory and frontal cortices (41, 42).

Multiple regression analysis included all time points from –30 to +200 ms and was conducted across all electrodes. Significant relationships with social and non-social touches were largely restricted to the electrodes above the contralateral sensorimotor cortex (contralateral to the stimulated hand), i.e., the electrodes that also showed the highest signal-to-noise ratio (***Figure 2c–f***). Our analysis revealed that the number of social touches was correlated with the thumb-associated signal-to-noise ratio at time points between 70 and 100 ms, and then again between 125 and 150 ms (***Figure 2c***). Notably and contrary to the simplistic prediction, the direction of the correlation was such that the higher the number of social touches, the lower the signal-to-noise ratio (***Figure 2c***). In contrast, the history of non-social touches was significantly related to the cortical signals in a narrow window between 135 and 150 ms, so that the higher the number of touches, the larger the signal-to-noise ratio (the relationships with other explanatory variables are presented in ***Figure 2 – Supplement 1*).**). These results suggested that social touches were tracked by the somatosensory cortex separately from non-social touches, and that the social touches were encoded at multiple stages of somatosensory processing.

To verify whether the uncovered relationship between the number of social touches on the phone and signal-to-noise ratio for the thumb was based on the social category per se, we once again employed random category shuffling. Based on the maximum signal-to-noise ratio, for the signal-to-noise ratio at the chosen electrode, the distribution of relationships for the number of touches on random categories was well separate from the relationship based on touches on Social Apps (***Figure 2g***). We also explored the relationships between the number of social touches on the phone and the somatosensory signal-to-noise ratios for the index and middle fingers, in addition to the thumb (***Figure 2h***). In comparison with the thumb, the relationships were substantially weaker for the index finger and absent for the middle finger. In summary, these results suggested that engaging in social activity on the touchscreen diminishes the cortical signal-to-noise ratio associated with the thumb, contrary to the anticipated consequences based on a simplistic view of use-dependent plasticity.

### Neuronal correlates of social touches on the keypad

The neuronal correlates of social touches described above were based on all touchscreen gestures, leaving open the possibility that the correlates reflected the underlying differences in the gestures used on Social vs. Non-social Apps. We matched the gesture type by restricting the analysis to pop-up keypads. A near-identical pattern of correlates was observed as in the original analysis that included all gestures. Briefly, with an increasing number of social touches on the keypad, the signal-to-noise ratio associated with the thumb between 70 and 100 ms decreased (***Supplementary Figure 4***).

### Social touches vs. somatosensory signal-to-noise ratio correlations as a function of time

According to the results presented above, the signal-to-noise ratio at early stages of the cortical somatosensory processing was significantly correlated with the number of social touches on the touchscreen but not with the number of non-social touches. Touchscreen behavior was continuously recorded prior to the EEG measurements. We leveraged this continuity to establish the temporal dynamics in terms of the time elapsed between the touchscreen behavior and the EEG measurement. Using the observations from the chosen electrode, we found the following complex temporal dynamics: the relationships were strong when examining recent social touches, followed by complex relationships decay, and the relationships picked up again with older touches (***Figure 2i***). The dynamics, although apparently more complicated than what was observed for the social touches vs. movement time variability relationships, were well captured using the following formula (R^2^ = 0.83):

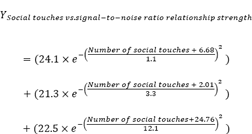

This relationship pattern suggested that a complex mix of both fast and slow mechanisms of plasticity is employed when configuring the cortex according to the history of social touches.

### Increased trial-to-trial variability in neuronal response amplitude is associated with social touches on the touchscreen

A reduction in somatosensory cortical signal-to-noise ratio associated with a larger number of social touches may be associated with two entirely different attributes of neuronal activity. First, the reduction may genuinely reflect an alteration in the amount of neuronal activity; and second, the reduction may reflect increased trial-to-trial temporal jitter, so that averaging of responses across trials results in a smaller amplitude (43). In theory, it would be possible to address these two possibilities by focusing on the shape of the evoked potentials at a single trial level to estimate the variability in peak amplitude separately from peak latency. However, in practice, the EEG signals intensely fluctuate at the single trial level, precluding facile analysis of the shape of the evoked potentials. To partly smooth the signals, we averaged a subset of 25 trials. Next, we detected the amplitude and latency of local maxima that immediately followed the temporal landmarks placed at 50 and 85 ms (***Figure 3a***). The landmarks were set so as to focus on the initial stages of somatosensory processing that did not encode the number of social touches according to the signal-to-noise ratio analysis (50 ms) and later stages that did (85 ms, at the center of the correlated range of 70–100 ms). We repeated this with a different subset of 25 trials, 10^5^ times for each volunteer, to estimate the trial-to-trial variability of the corresponding latencies and amplitudes (***Figure 3b,c***).

**Figure 3.**
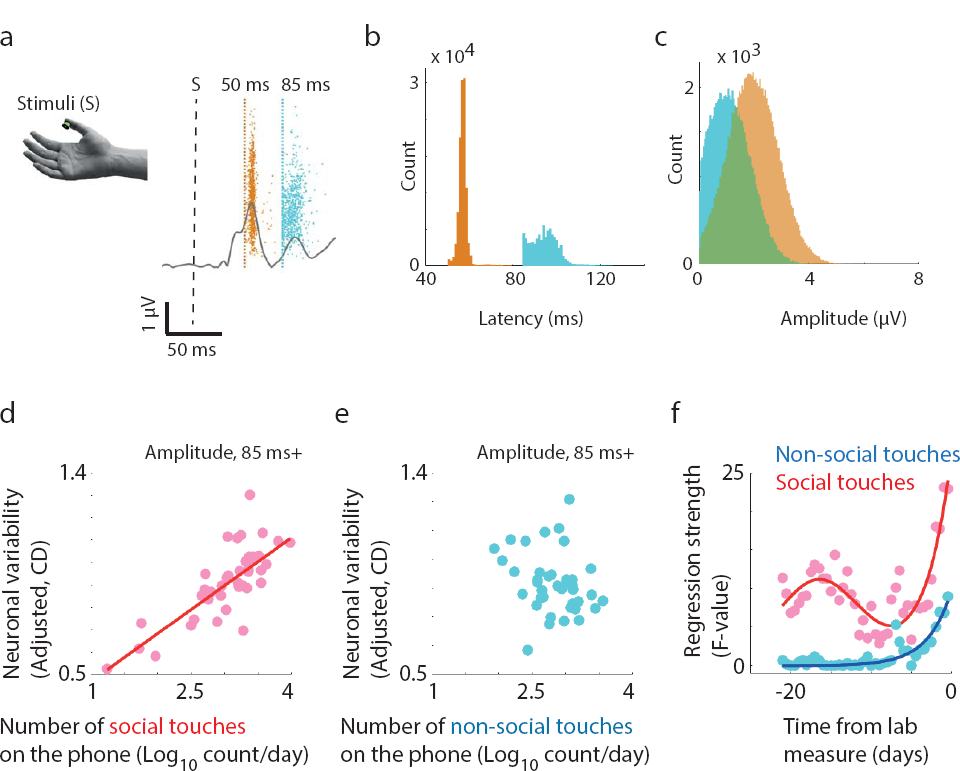
The trial-to-trial variability in the degree of cortical responses is proportional to Social App usage. (**a–c**) Depiction of the analysis method to separately estimate the trial-to-trial variability in the cortical signal latency and the amplitude. (**a**) Rectified event related potentials based on a random sample of 25 trials was generated 10^5^ times. The rectified potential based on all the trials in one volunteer is drawn in grey. The first local maxima encountered on 10^3^ iterated potentials after the set temporal landmarks of 50 and 85 ms are indicated (colored dots). The distribution of latencies (**b**) and amplitudes (**c**) of the first maxima in the same volunteer based on which the corresponding coefficient of dispersion (CD) was estimated. (**de**) Adjusted response plots. (**d**) The greater the number of social touches in the 21-d recording period, the larger the variability in signal amplitudes at the 85 ms landmark (measured in terms of CD). (**e**) The relationship between the number of non-social touches and the variability was not significant. (**f**) Parsing the touchscreen recordings in 12 h steps (72 h bin) revealed that the relationship for non-social touches simply decayed with older touchscreen data and a more complex pattern was apparent for the social touches.

The variability of cortical signal amplitudes detected by the 50 ms landmark was unrelated to the explanatory variables that included movement time variability in addition to the original set of variables derived from the touchscreen and gender [R^2^ = 0.31, *f* (7,33) = 2.11, *p* = 0.07, robust linear regression]. In particular, amplitude variability was clearly unrelated to the number of social touches [*t*(1,33) = 0.68, *p* = 0.5] and non-social touches [*t*(1,33) = −0.02, *p* = 0.98, ***Supplementary Figure 5***]. The variability of signal latencies at this temporal landmark was also unrelated to the social touches [*t*(1,33) = 0.60, *p* = 0.6] and non-social touches [*t*(1,33) = −0.23, *p* = 0.8, ***Supplementary Figure 5***]. In contrast, the variability of signal amplitudes detected by the 85 ms landmark was strongly related to the explanatory variables [R^2^ = 0.45, *f* (7,33) = 3.9, *p* = 0.003, robust linear regression]. We observed that the higher the number of social touches, the larger the variability [*t*(1,33) = 4.62, *p* = 5.6 × 10^−5^, ***Figure 3d***]. There was a weak trend linking the number of non-social touches and neuronal variability, such that the higher the number, the lower the variability [*t*(1,33) = −1.9, *p* = 0.07, ***Figure 3e***]. In terms of variability of signal latencies at this landmark, a weak relationship with the explanatory variables was observed [R^2^ = 0.34, *f* (7,33) = 2.5, *p* = 0.04, robust linear regression], and the higher the number of social touches, the larger the neuronal temporal variability [*t*(1,33) = 2.3, *p* = 0.03, ***Supplementary Figure 5***]. Finally, we did not find any significant links between movement time variability and neuronal response variability [latency dispersion at 85 ms: *t*(33) = −1.8, *p* = 0.08; amplitude dispersion at 85 ms: *t*(33) = −1.91, *p* = 0.06]. This raised the possibility that although both movement time variability and neuronal variability increased with social touches, the two measures themselves reflected largely separate neuronal process.

In summary, the results were consistent with the notion that trial-to-trial variability of both, the degree and timing of neuronal activity, increased according to the number of social touches. However, it must be noted that the evidence for increased temporal variability was rather weak in contrast with the evidence for increased amplitude variability.

### Time-dependent structure of the relationships between touchscreen use and neuronal variability

As with the preceding time-dependent analyses, we reasoned that the putative plasticity attributes could be studied by sampling touchscreen behavior at various times before laboratory measurements. Since a tendency was observed linking non-social touches over the entire recording period with neuronal variability, we first studied temporal dynamics of the phenomenon using F-values associated with non-social touches. The relationship strength simply decayed as a function of time and was well described by the following formula (R^2^ = *0.*81*, **Figure 3f**):*

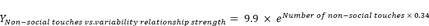

The social touches showed more complex dynamics, such that the relationship was strong when using recent touchscreen data, weakening over time. The relationship was also strong when using older data. This was well captured by the following equation (R^2^ = 0.72, ***Figure 3f***):

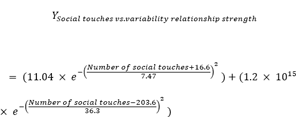

Time-dependent neuronal variability dynamics of the correlates were qualitatively similar to what we observed for motor time variability. Overall, these results indicated that social touches are distinctly integrated to reconfigure the cortical circuits associated with the thumb and both rapid and slow forms of use-dependent plasticity are employed towards this putative reconfiguration.

## Discussion

One striking finding of this report was that the individuals who generated a larger number of social touches on the touchscreen were more variable in their response times when performing a simple task with the thumb. The somatosensory cortical activity also exhibited more variability associated with social touches. The dense digitization of behavior on the smartphone allowed us to quantify and contrast these relationships with the history of non-social touches. The results based on social touches data were contrary to the simplistic view of use-dependent plasticity, which predicted more stable sensorimotor computations corresponding to an increased touchscreen use. Even when placed outwith the framework of use-dependent plasticity, these results suggested that the understanding of inter-individual differences in elementary sensorimotor control is deeply inter-connected with the details of behavior expressed in the real world.

We interpret these results as indicative that social activities on the touchscreen lead to increased sensorimotor variability. However, the correlational nature of our findings precludes us from discarding an alternative possibility that a higher sensorimotor variability leads to more social touches, or that a common factor determines both these variables. Based on the current knowledge, a reasonable case for the former cannot be made but the latter must be seriously considered. Extraverted individuals are characterized by higher usage of Social Apps than introverts and extraversion is associated with diminished somatosensory cortical activity evoked by the fingertips (44, 45). The extraversion-based relationship is specific to the left hand and is absent for the right hand (45). In contrast, our study focused on the right hand. Moreover, the extraversion-based relationship is not specific to particular fingertips, in contrast to the thumb-specific correlates of touchscreen use uncovered here and in our previous study (16). In addition to the personality factor, cognitive states that lead to enhanced attention or arousal may influence both the touchscreen behavior and neuronal measures in the laboratory (46). This state-dependent view does not account for the observation that touchscreen-based correlates were largely restricted to the thumb. It also does not account for how the 1-2 weeks old touchscreen data could strongly correlate with the laboratory measurements. Given these evidences, the framework of use-dependent plasticity may be the most appropriate for considering our findings.

Neuronal correlates uncovered here suggest that low-level sensorimotor processing, at the primary somatosensory cortex, encodes the history of social touches on the touchscreen. This observation is consistent with the notion that the primary sensory areas do not exclusively represent the incoming sensory inputs but integrate these inputs into behavioral context (47). For the somatosensory cortex, this is supported by laboratory observations that the cortex participates in multi-sensory integration and that factors, such as attention, modulate its activity and plasticity (4, 48, 49). Our findings provide a real-world example that the behavioral context of an experience is a key factor in configuring the cortex.

The temporal dynamics of the associations uncovered herein provide some insights into the nature of processes engaged in the putative use-dependent plasticity. For both, trial-to-trial movement time variability and neuronal variability, we observed a complex fall and then rise in the relationships strength with older data from the Social Apps. This pattern suggests that social touches trigger both rapid and slow mechanisms of plasticity. Rapid mechanisms may include such processes as alteration in excitatory-inhibitory balance or the unmasking of preexisting circuits (8, 50). Slow mechanisms may include the formation of entirely new pathways, comprising changes of the underlying white matter that may take weeks to complete (5, 51). The relationship with older data from the Non-social Apps simply decayed, suggesting exclusive deployment of rapid mechanisms.

It is not clear how the sensorimotor cortex sorts the touches on Social Apps separately from Non-social Apps. One possibility is that the social touches are sorted based on top-down information flow via neuromodulators or feedback from high-level neuronal networks engaged in social behavior (14, 51). Another possibility is that the touches are sorted in a bottom-up manner based on distinct sensory features that accompany the social touches. We tested this possibility by restricting our analysis to pop-keypad touches, only to discover that even when the gestures were apparently matched, the social touches showed a distinct sensorimotor correlate. Other relevant but unexplored differences in the input statistics of Social vs. Non-social Apps may exist in terms of the length of the words typed or the complexity of language used. Nevertheless, a previous study on typing skills suggested that greater experience was associated with smaller sensorimotor variability (23). Therefore, the increased variability associated with social touches cannot be easily explained using the widely held notions on use-dependent plasticity.

Why does sensorimotor variability increase with social touches on the touchscreen? We propose that the increased variability is an inevitable consequence of repeated engagement of the thumb in social cognition. Essentially, social touches on the touchscreen are accompanied by an array of neuronal processes associated with language, anticipation, and social status (13). Presumably, using Hebbian-like mechanisms of plasticity, the thumb becomes increasingly connected with this broad array of processes. It is this enhanced embedding of sensorimotor processing in a broad array of neuronal processes that may lead to increased noise in low-level circuits (53).

In the population of young adults sampled here, the median number of touchscreen touches generated per day was 2.7 × 10^3^ and the most active individual generated 1.1 × 10^4^ touches per day. These numbers reflect the dominance of touchscreen events in modern human actions, comparable in magnitude with the number of steps (1 × 10^4^) or eye blinks per day (1.2 × 10^4^) (54, 55). Therefore, it should not be surprising that the neuronal sensorimotor processing is reconfigured by touchscreen behavior (16). The nature of the touchscreen behavior-neuronal relationships uncovered by leveraging seamless quantifications on the smartphone warrants a more in-depth examination on how social activities on the touchscreen reconfigure the brain. These links also highlight the complex nature of neurobehavioral relationships in elementary sensorimotor control, such that the history of social and non-social touches, the rate of touchscreen activity, and number of different Apps used are all independently encoded to impact future computations. Addressing how the quantitative history of touchscreen behavior relates to elementary neuronal functions will help bridge the large gap between inherently artificial laboratory experiments and the behavior expressed in the real world.

## Materials and Methods

### Subjects

Volunteers (n = 57) were recruited using campus-wide announcements at the University of Zurich and ETH Zurich between December 2014 and August 2015. The sample consisted of subjects within a narrow age group [26 females; 23 (20th percentile) to 28 (80th percentile) years old]. The age at which the volunteers reportedly began using the phone was also narrowly distributed [19 (20th percentile) to 25 (80th percentile) years old]. Previous reports on inter-individual variability in cortical somatosensory signal-to-noise ratio, touchscreen use-dependent plasticity and use-dependent reduction in sensorimotor variability employed a sample size between 15 – 28 (16, 18, 23, 55). Essentially we anticipated a weaker impact of the social touches on the touchscreen than the explanatory variables studied before, i.e., deliberate laboratory practice, touchscreen use in general and the presence of autism spectrum disorder. Therefore, we doubled the sample size and employed more regression parameters than the previous studies to increase the sensitivity of our analysis. All experimental procedures were conducted according to the Swiss Human Research Act approved by the cantons of Zurich and Vaud. The procedures also conformed to the Declaration of Helsinki. The volunteers provided written and informed consent before participating in the study. Reasonable health, right-handedness, and ownership of a non-shared touchscreen smartphone were pre-requisites for participation. The handedness was further verified by a questionnaire (57). The fingers used on the touchscreen were analyzed using a pictorial survey where the volunteers ranked each finger on a scale 1–10 (1, least preferred; 10, most preferred).

### Smartphone data collection and analysis

A custom-designed background App was installed on the volunteers’ smartphones to quantify the touchscreen behavior (see the Supplementary Methods for in-depth description of the design and performance specifications of the App). Briefly, the App recorded the timestamps of touchscreen events and the label of the App on the foreground. The App recorded the touchscreen events with an interquartile error range of 5 ms. Data were stored locally and transmitted by the user at the end of the observation period via secure email. Smartphone data were processed using custom written scripts on MATLAB (MathWorks, Natik, USA). In smartphones with more relaxed permission settings (build-in), the pop-up keypad touches were additionally labeled. The number of touches on each App category (“Social”, “Non-social”, or “Uncategorized”) was divided by the length of the recording period to determine the number of touches per day. Apps that functioned to enable social interactions between a circle of friends or acquaintances were labeled as “Social” and Apps that clearly did not feature this functionality were labeled as “Non-social”. Apps whose label was poorly registered by the operating system, untraceable on Google Play, or that contained both social and non-social properties, e.g., gaming Apps with social messaging, were labeled as “Uncategorized”. The touches that were separated by less than 50 ms were eliminated from further analysis. The rate of touchscreen events was determined as 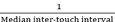. A recording period of up to 21 d was used for the main regression analysis. The number of Apps that were used over the recording period was counted.

### Simple reaction time task and analysis

Volunteers responded to a brief (10 ms) tactile pulse by depressing and releasing a button mounted on a micro switch. The tactile pulse was presented by using a computer-controlled solenoid tactile stimulator (Heijo Research Electronics, London, UK). The stimulating magnetic rod (2 mm in diameter) generated a supra-threshold 2-mN contact. The thumb or the middle finger was stimulated. The micro switch (extracted from RX-300 optical mouse, Logitech, Lausanne, Switzerland) was operated by press-downwards and release-upwards movements of the thumb or the middle finger. All volunteers performed the task with the thumb (n = 57) and a subset of randomly chosen volunteers performed the task with the middle finger in addition to the thumb (n = 17).

The task was repeated 500 times (for each fingertip) within an experimental session, with 2 min break in the middle of the session. The pulses were delivered with 3 ± 1 s gap and the button presses generated analogue signals that were digitized at 1 kHz. The reaction time and movement time (the time taken to execute button depression) were fitted with three ex-Gaussian parameters. This form of fitting separates skewed reaction time data into a Gaussian region and an exponential region. Mean of the Gaussian region was captured by parameter μ, and variability of the Gaussian region by parameter σ. The exponent τ captured unusually slow responses. The parameters were estimated using previously described MATLAB scripts (36).

### EEG data acquisition and analysis

A subset of volunteers (n = 43) participated in EEG experiments. The volunteers were seated upright for the EEG and the right, stimulated, hand was concealed by a baffle. Computer-controlled solenoid tactile stimulator (see above) was attached to the right thumb tip and to the right index and middle finger tips. To ease the tedium of the hours-long measurements required for gathering the tactile evoked potentials data (SSEPs), volunteers were allowed to view a movie (David Attenborough’s Africa series); white noise, played to mask the sound generated by the stimulator, was mixed with the movie soundtrack and delivered through headphones. The number of trials was set to 1000 for each fingertip, randomized for the tips, and the stimuli were separated for each fingertip by 2–4 s. A non-alcoholic and caffeine-free drink break was offered every 10 min, for a maximum of 10 min. To record the EEG signals, 64 electrodes were used (62 equidistant scalp electrodes and two ocular ones), against a vertex reference (EasyCap, Herrsching, Germany), as previously reported (16). The electrode locations were digitized in a 3D nasion-ear coordinate frame (ANT Neuro and Xensor software, Netherlands) for a representative volunteer. The signals were recorded and digitized by BrainAmp (Brain Products GmbH, Gilching, Germany) at 1 kHz. Offline data processing was accomplished using EEGLAB, a toolbox designed for EEG analysis on MATLAB (58). The data were referenced to the average of all scalp electrodes and band-pass filtered between 1 and 80 Hz. “Epoched” trials over 80 μV were eliminated to remove large signal fluctuations, e.g., from eye blinks. The data were further processed using independent component analysis. Components dominated by eye movements and other measurement artifacts were eliminated by using the EEGLAB plug-in SASICA (59). The signal-to-noise ratio was estimated using the linear modeling toolbox LIMO EEG (EEGLAB plug-in) (60). In this toolbox, R^2^ values were estimated for each volunteer based on single trials, as a sum of squares of the putative signal divided by the sum of squares of the residuals. Essentially, the predominant notion in the sensory evoked potential research field is that the average over multiple trials extracts a signal that is otherwise hidden in the measurement noise and background neuronal processes (39). The signal-to-noise ratio in this case captures how well the estimated mean (putative signal) represents the data. To normalize the data across the sampled population, the square root of the putative signal-to-noise ratio was used for subsequent analyses using multiple linear regression.

The trial-to-trial variations in EEG responses were estimated based on the rectified event-related waveforms of 25 randomly sampled samples. The resampling was reiterated 10^5^ times for each individual. The first local maxima above 50 and 85 ms were estimated for each iteration. The maxima were estimated using a MATLAB add-on function (“EXTREMA”). This form of bootstrapping was used to recover the distribution of signal timings and amplitudes, and these distributions were subsequently used to derive the coefficient of dispersion for each individual 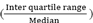 at marked time points.

### Correlational statistics

All analyses involving the reaction and movement times were conducted by robust-bi-square-multiple linear regression analysis (implemented in MATLAB). The fitted model was evaluated using ANOVA with a level of significance set at *p* = 0.05. The following main regression equation was used:

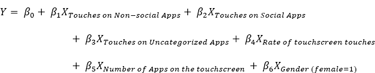

Where *Y* took the form of ***Y_Movement time variability_*** or *Y_Reaction time variability_*, or *Y_Somatosensory putative signal-to-noise ratio._* For *Y_Coefficient of dispersion in peak latency_* and *Y_coefficient of dispersion in peak amplitude_*, the explanatory variable *β*_7_*X_Movement time variability_* was added to the original equation. *β*_1 to *n*_ comprised regression coefficients estimated by robust regression, and *β*_0_the intercept. The explanatory variables quantifying the touchscreen behavior were based on 21 d of recording made prior to the laboratory measures.

To analyze the time-dependent structure of regression parameters associated with the number of touchscreen touches, we used the following approach. The parameters *X_Touches on Non-social Apps_*, *X_Touches on Social Apps_*, and *X_Touches on Uncategorized Apps_* were re-estimated over the span of 21 d with 12-h steps and 72-h windows. Other parameters were unchanged and, as in the main regression equation, were based on the data spanning the entire 21-d period. To describe the time-dependent fluctuation of F-values, the relationship was iteratively fitted by comparing linear, exponential, and Gaussian equations with a maximum of three terms. The fit with the highest R^2^ was used to describe the relationships.

Similarly, to assess the temporal structure of the variable typical rate of touchscreen use or the number of Apps used, the variables *X_Rate_* or *X_Number of Apps on the touch screen_* were re-estimated with 12-h steps and 72-h windows while other parameters remained unchanged.

As a control, we repeated the analysis with shuffled App categories. Essentially, for the original analysis, the Apps were labeled as “Social”, “Non-social”, and “Uncategorized” according to a fixed criterion, i.e., Social Apps were those that enabled the communication of a message or an opinion to a circle of friends or acquaintances. The list of all Apps in the database and their classifications were randomly shuffled (10^5^ iterations). These shuffled lists were then used to estimate the number of touches in each of the action categories. Note that the total number of Apps in each category was constant during shuffling.

Plots for displaying multiple linear regression results in two dimensions (adjusted response plots) were generated using a built-in MATLAB function (plotAdjustedResponse). Formulation of this plotting method and its advantages are described elsewhere (61).

The EEG data were correlated with touchscreen parameters using robust regression, iterative least squares method (implemented in LIMO EEG). The correlation coefficients were estimated across all electrodes and for the time period from -30 to 200 ms relative to the stimulation onset. When focusing the analysis on keypad use, due to the smaller number of samples, the variables were restricted to parameters *X_Rate_*, *X_Number of touches on Social Apps_*, and *X_Number of touches on Non-Social Apps_*. The regression statistics were corrected for multiple comparisons by using 1000 bootstraps and spatiotemporal clustering, as implemented in LIMO EEG.

## Acknowledgements

The data collection was made possible by the assistance of Ciara Shortiss and Magali Chytiris. We thank Enea Ceolini for helping in the design and implementation of the behavioral tracking software. This research was funded by Holcim Stiftung and the Society in Science Branco Weiss Fellowship. The authors would like to thank Eric Rouillier, Anne-Dominique Gindrat, Kevan Martin for discussions. We are grateful to Valerio Mante for making the crucial suggestion of randomizing the App categories. The authors thank Joanna Mackie for help in editing this manuscript.

## Author Contributions

MB acquired the data, participated in data analysis, and edited this manuscript. AG designed the study, helped in data acquisition, analyzed the data, and drafted this manuscript.

## Supplementary Information Index

**Figure 1 - Supplement 1.**
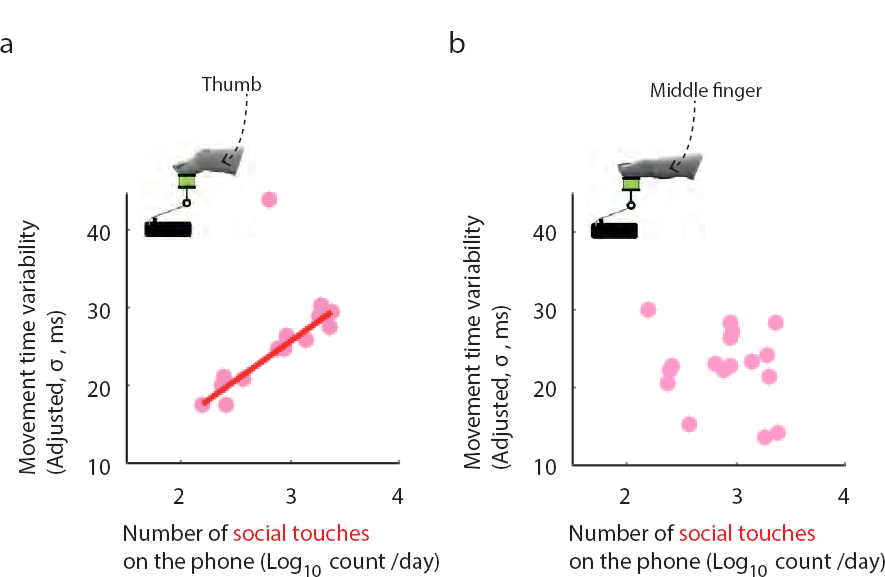
The social touches do not reflect on movement time variability when the task is performed with the middle finger. (a) Adjusted response plot showing the link between the number of social touches generated on the touchscreen and the movement-time variability when the task was performed by using the thumb. Specifically, higher the number of social touches the higher the movement time variability (b) When the same volunteers performed the task with the middle finger the relationship was absent.

**Figure 1 - Supplement 2.**
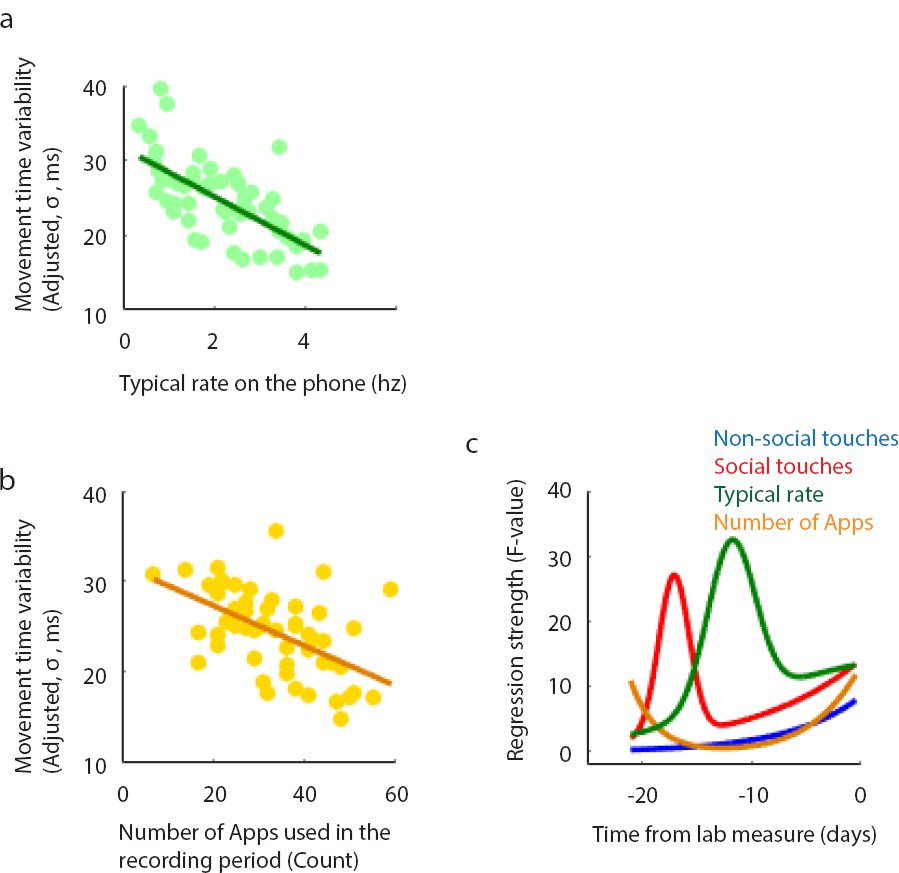
Analysis of explanatory variables other than the number of social and non-social touches. (a-b) Adjusted response plots. (a) The link between the typical rate of touchscreen usage and movement time variability and (b) the number of Apps used and the variability. (c) The analysis of the relationships to movement time variability after parsing the touchscreen recordings in 12 h steps (72 h bin).

**Figure 2 - Supplement 1.**
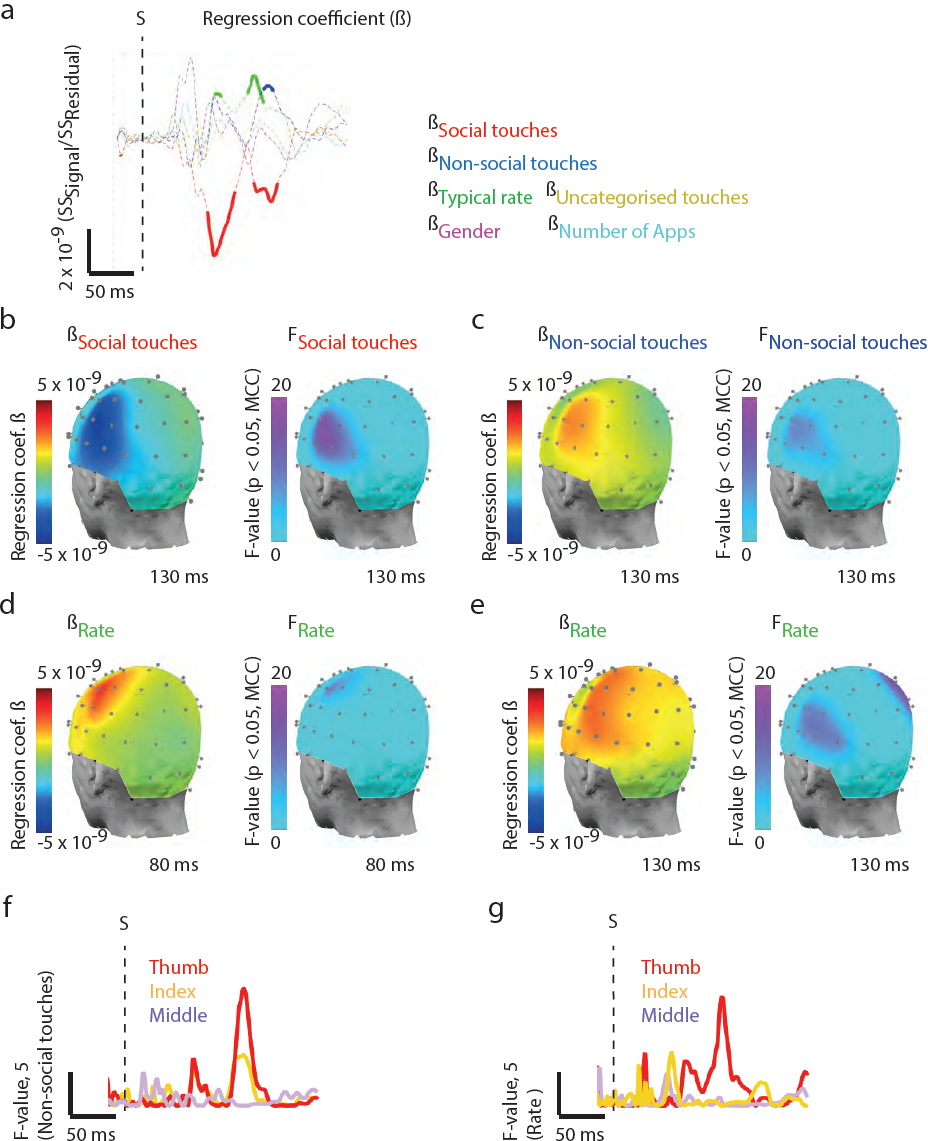
The trial-to-trial variability in the degree of cortical responses is proportional to Social App usage. **(a-c)** Depiction of the analysis method to separately estimate the trial-to trial variability in the cortical signal latency and the amplitude. **(a)** Rectified event related potentials based on a random sample of 25 trials was generated 105 times. The rectified potential based on all the trials in one volunteer is drawn in grey. The first local maxima encountered on 103 iterated potentials after the set temporal landmarks of 50 and 85 ms are indicated (colored dots). The distribution of latencies **(b)** and amplitudes **(c)** of the first maxima in the same volunteer based on which the corresponding coefficient of dispersion (CD) was estimated. **(d-e)** Adjusted response plots. **(d)** The greater the number of social touches in the 21-d recording period, the larger the variability in signal amplitudes at the 85 ms landmark (measured in terms of CD). **(e)** The relationship between the number of non-social touches and the variability was not significant. **(f)** Parsing the touchscreen recordings in 12 h steps (72 h bin) revealed that the relationship for non-social touches simply decayed with older touchscreen data and a more complex pattern was apparent for the social touches.

Supplementary Methods: Description of the App used to track touchscreen behavior.

Supplementary List: A sample of all the Apps in the database to illustrate the App categorization used in this study in Social and Non-social Apps.

Supplementary Figure 1: The plot matrix of the explanatory variables and the corresponding variation inflation factors.

Supplementary Figure 2: Social touches on the keypad is related to movement time variability. (a-b) Adjusted response plots. (a) Higher the number of social touches on the touchscreen pop up keypad the higher the movement time variability. (b) The non-social touches on the keypad were not related to the variability.

Supplementary Figure 3: The reaction time variability is related to the number of social touches. (a) Adjusted response plot displaying that higher the number of social touches the larger was the reaction time variability. (b) The non-social touches were unrelated to the reaction time variability. (c) The relationship discovered for the social touches was well apart from the distribution of relationships obtained by using randomly shuffled categories.

Supplementary Figure 4: The neuronal correlates of the number of social touches on the touchscreen keypad. When we restricted our analysis to the pop-up keypad touches, we found that higher the number of social touches on the keypad smaller the signal-to-noise ratio as in the original analysis including all types of touchscreen events. The legend is identical to Figure 2 panels a-f.

Supplementary Figure 5: The neuronal variability determined from the early temporal landmark set at 50 ms was unrelated to the number of touches. (a-d) Data by using the 50 ms temporal884 landmark. Adjusted response plots showing the non-significant regressions between social or non-social touches and neuronal variability in terms of amplitude or latency. (e,f) Latency data by using the 85 ms temporal landmark shows a weak relationship between social touches (and not for non-social touches) and trial-to-trial temporal variability.

